# A ceramide synthase is important for filamentous fungal biofilm morphology and antifungal drug resistance

**DOI:** 10.1101/2025.11.16.688756

**Authors:** Charles T.S. Puerner, Owen M. Wilkins, Robert A. Cramer

## Abstract

The complex structure of fungal biofilms generates microenvironments that impact the fitness of cells within the biofilm community. Contributions to fitness include the development of emergent properties resulting in the tolerance or resistance to external stressors such as rapid environmental changes and in the context of an infection, antifungal drug exposure. The biofilm developed by the filamentous fungal pathogen *Aspergillus fumigatus* develops zones of low oxygen which contribute to a reduction in antifungal drug susceptibility, however the genes and mechanisms involved in driving this emergent property of the biofilm are ill-defined. In this study, we utilized a transcriptomic approach to probe the biofilm structure in comparison to drug susceptible planktonic cultures to identify transcriptional patterns and genes unique to the *A. fumigatus* biofilm. Importantly we utilized two phenotypically diverse strains that allowed us to identify biofilm specific gene co-expression networks. One of these networks was highlighted by a gene encoding a ceramide synthase, designated *barA*, with a striking increase in *barA* transcript abundance specifically in the biofilm. Null mutants for *barA* in two strain backgrounds display altered biofilm morphology with some strain specific differences. Importantly, *barA* has a role in regulating susceptibility to ergosterol targeting antifungal drugs. These data identify biofilm specific genes in *A. fumigatus* for further study and highlight the importance of fungal ceramide synthases in mediating antifungal drug susceptibility in infection relevant biofilms.

**Importance:** Biofilms are problematic structures in the context of microbial infections due to the ability to resist both host and drug mediated attempts at tissue sterilization. Consequently, it is imperative to identify mechanisms underlying the development of these structures and the emergent properties they develop. The filamentous fungal pathogen *Aspergillus fumigatus* forms robust structured biofilms that are resistant to contemporary antifungal drug treatments though the mechanisms are ill-defined. In this study we compared the transcriptional landscape of *two A.* fumigatus reference strains grown as biofilms and in planktonic culture conditions to identify biofilm specifc genes and pathways. These analyses and subsequent genetic and phenotype studies revealed that a ceramide synthase is important for the development of antifungal drug resistant biofilms. Consequently, these data support the rationale for targeting fungal lipid homeostasis for antifungal therapeutic development, particularly in the context of biofilm mediated infections.

## Introduction

Microbial biofilms form complex microenvironments that require dynamic spatial and temporal cellular responses (1–3). Biofilms contribute to virulence and are important for many organisms’ pathogenicity (4). In contrast to often studied homogenous and stable laboratory planktonic or batch cultures, biofilms represent an opportunity to investigate a heterogenous microbial community that is temporally dynamic with shifting internal environments and stresses. The cellular adaptations of cells within the biofilm to these dynamic microenvironmental changes contribute to emergent properties of the biofilm such as antimicrobial resistance or resistance to host defense systems (5–7).

Filamentous fungi form networks of hyphae, termed a mycelium, with emergent properties similar to well-studied bacterial biofilms (8–16). Studies on filamentous fungal biofilms are emerging, particularly in the human pathogen *Aspergillus fumigatus*, a normally saprophytic mould that is found ubiquitously in the environment (17, 18). In the context of human health *A. fumigatus* is capable of causing severe pulmonary infections in immunocompromised individuals where inhaled conidia often form biofilm like structures in tissue (19, 20). These fungal biofilms are difficult for the host immune system to overcome and difficult to treat with antifungal therapies (5, 21). Taken together it is important to understand the physiology of *A. fumigatus* in the context of the biofilm as this closely resembles the organism at the site of an established infection.

*In vitro A. fumigatus* submerged culture biofilms form morphologically distinct structures that are similar to structures observed within the infection environment (8, 9). Key features of the *in vitro* biofilm model that replicate *in vivo A. fumigatus* growth is a lack of asexual development structures and production of an extracellular matrix composed largely of galactosaminogalactan polymers. *A. fumigatus* biofilms also form steep low oxygen gradients that have been shown to drive the emergent property of increased antifungal resistance and are also observed at the site of infection (9). The mechanism driving this antifungal drug resistance is ill-defined. It has been suggested that extracellular DNA (eDNA) accumulation and efflux pump activity in the biofilm are also involved, but significant questions remain regarding the genes involved and underlying mechanisms (11, 12).

As much of our understanding of *A. fumigatus* at the transcriptional level has come from studies utilizing planktonic cultures or cultures on solid medium that rapidly undergo asexual development (colony biofilms) there is a lack of understanding of the transcriptional landscape of the submerged biofilm (8, 22–26). In this present study we observe through transcriptional profiling of submerged culture biofilms and planktonic cultures of two reference *A. fumigatus* strains in two distinct oxygen conditions that the biofilm transcriptional landscape is unique. Utilizing these data, a ceremide synthase was discovered to be a mediator of biofilm formation and antifungal susceptibility. Consequently, these data suggest inhibition of fungal ceramide synthesis is a promising anti-biofilm strategy for future therapeutic development.

## Results

### *Aspergillus fumigatus* biofilm transcriptional landscape is distinct from batch culture

Given the relevance of the *A. fumigatus* biofilm to the growth state in the host environment and its unique properties, we sought to define genes specific to the biofilm growth state. We were also curious if the more well studied planktonic culture hypoxia response would be similar to the biofilm state given the oxygen gradients previously observed in *A. fumigatus* biofilms (9). We utilized RNA-Seq based transcriptional profiling of 18 hour submerged biofilms and planktonic cultures with and without exposure to low oxygen (0.2% O_2_) for 30 minutes to induce a hypoxia response in two diverse reference strains (CEA10/A1163 and AF293) (**Fig. S1A).**

Global Euclidean distance clustering reveals oxygen tension as a major determinant of hierarchical clustering (**Fig. S1B**). Unexpectedly, the biofilm samples more closely clustered with the normoxia planktonic samples indicating that the response to low oxygen in a biofilm is distinct from planktonic culture hypoxia response. There was additional clustering of samples by growth condition and strain indicating the biofilm is a unique state from both the low oxygen and normoxia samples. Using a principal component analysis of the top 4000 most variable genes, we found that 79.6% of the variability in the data was explained by the first three principal components (**Fig. S1C-F**). A 4000 gene threshold was set as this encompasses the majority of variability within the data (**Fig. S1G**). Principal component 1 appears to be explained by oxygen tension. Principal component 3 separates the biofilm samples from the planktonic samples indicating there is a subset of the data that is specific to the biofilm state. Cluster analysis of the top 4000 most variable genes revealed a distinct clustering pattern for each growth state with the biofilm appearing to be an intermediate between the low oxygen and atmospheric oxygen planktonic cultures but still clustering with the normoxia samples (**Fig. S1H**). From these analyses we found the biofilm state to be transcriptionally unique and therefore confident that biofilm specific transcriptional pattern and genes could be defined.

### Differential expression analysis reveals biofilm specific expression patterns

A differential expression analysis comparing the biofilm state to the planktonic state (both low oxygen and normoxia samples) identified a total of 1607 genes with increased abundance and 1591 genes with decreased transcript abundance in the biofilm samples (adjusted p-value < 0.05) (**Fig. 1A,B, Tables S1,S2**). A gene set enrichment analysis (**Table S3**) using FunCat, GO, and KEGG terms of the significant differentially expressed genes indicates the biofilm is distinct from the planktonic state in part through differences in metabolism, in an altered cell cycle, and altered gene expression regulation (**Table 1)**. Using an adjusted p-value cutoff of less than 0.05 the GSEA analysis idenfied 61 significant FunCat terms, 64 significant GO terms, and 11 significant KEGG terms. Among the genes with increased mRNA abundance in the biofilm compared to the planktonic condition there is a significant enrichment for fatty acid metabolism, oxidoreductase, FAD binding, and nitrogen metabolism genes. For example, regarding differences in metabolism in the biofilm among the genes with significant increase in transcript abundance is the putative acetate kinase gene Afu3g10750 and the isocitrate lyase *acuD* (**Fig. S1I)**. This indicates a potential shift in biofilm metabolism by generating acetate and acetyl-coA through fermentation for shuttling through the glyoxylate shunt. Additionally, there is an enrichment for siderophore transport in the biofilm indicating potential iron limitation possibly due to iron requirements for biofilm specific metabolism such as related to a biofilm specific hypoxia response. Among the genes with decreased mRNA abundance in the biofilm compared to planktonic conditions there is an enrichment for genes involved in cellular growth, cell cycle and gene expression regulation strongly indicating the cells within the biofilms are in an altered cell cycle state (**Table 1**). For example, the cell cycle regulator gene *cdc48* has a decrease in transcript abundance in the biofilm as well as the genes for the septin *aspA*, actin, and beta-tubulin indicating at least a subpopulation in the biofilm is likely not undergoing active growth (**Fig. S1I**). These data suggest the biofilm differs from the homogenous planktonic state (normoxic and hypoxic) by undergoing shifts in metabolic pathways needed for cell survival and maintenance of the biofilm.

**Figure 1:**
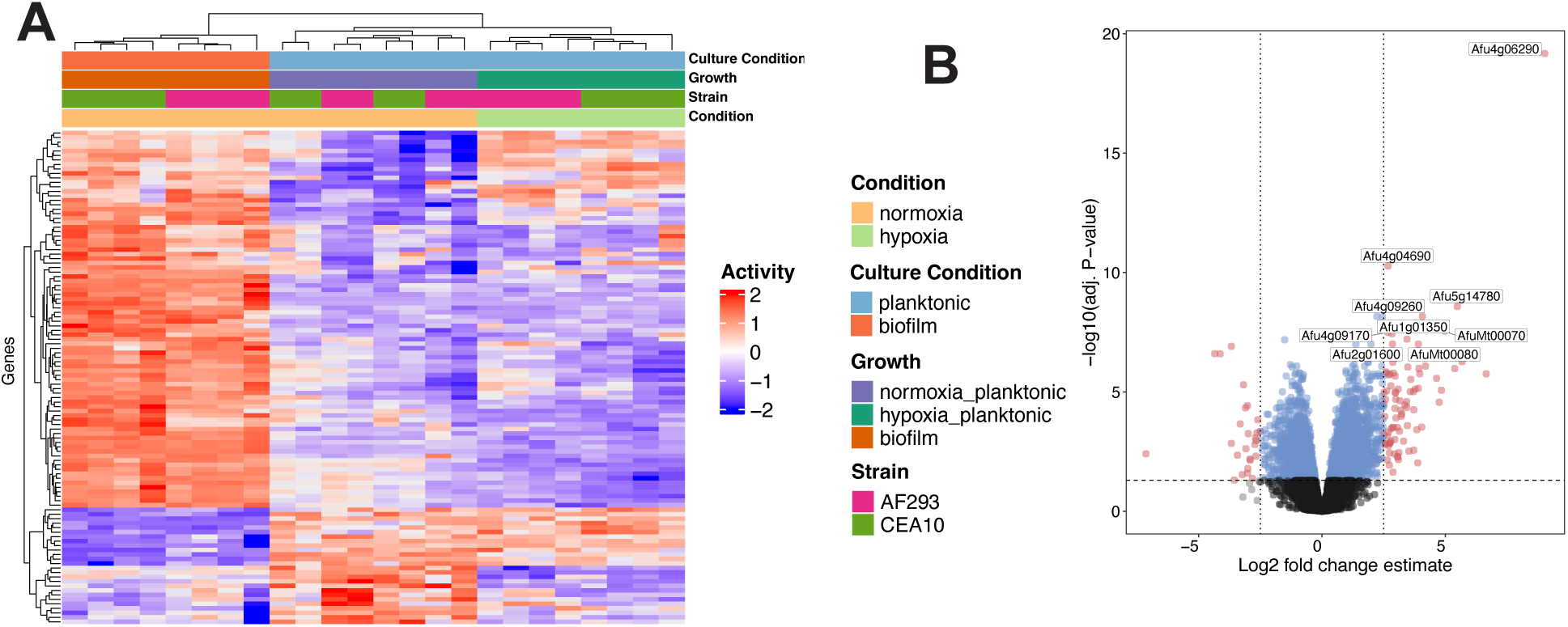
The *Aspergilus fumigatus* biofilm has a unique transcriptional state. Differential gene expression comparing biofilm samples to planktonic samples reveals biofilm specific genes. **A)** A heatmap representing genes with an absolute log2 fold-change of at least 2.5 and an adjusted p-value of less than 0.05. Unsupervised clustering confirms these are biofilm specific expression patterns. **B)** A volcano plot highlighting the significant DEG with an absolute log2 fold-change of at least 2.5 in red.

**Table 1:** Summary of GSEA of significantly differentially expressed genes using three functional category enrichments.

| Functional Theme | Status | Key Ontology Terms |
| --- | --- | --- |
| Metabolic shifts & transport | Upregulated | **FunCat:** siderophore-iron transport, fatty acid metabolism (NES +)<br>**GO:** xenobiotic transporter, oxidoreductase, FAD binding (NES +)<br>**KEGG:** nitrogen metabolism, taurine/hypotaurine metabolism (NES +) |
| Cytoskeleton & cell morphology | Downregulated | **FunCat + GO:** actin cytoskeleton, microtubule cytoskeleton, cell polarity, hyphal growth (NES -)<br>**KEGG:** motor proteins (NES -) |
| Cell cycle & division | Strongly downregulated | **FunCat + GO:** cell cycle checkpoints, mitotic cycle, spindle pole body, cytokinesis (NES -)<br>**KEGG:** cell cycle - yeast (NES -) |
| Gene expression machinery | Strongly downregulated | **FunCat + GO:** transcriptional control, RNA processing, ribosome biogenesis, RNA binding, DNA binding, chromatin organization, DNA replication (NES -)<br>**KEGG:** RNA polymerase, ribosome biogenesis, nucleocytoplasmic transport, cell cycle (NES -) |

### A weighted gene co-expression network analysis (WGCNA) reveals biofilm specific gene modules

A WGCNA was used to leverage the complexity of this dataset in order to identify modules of genes with correlated mRNA abundance profiles (27, 28). Using a soft threshold of 20 the constructed network achieved a scale free topology with an R^2^ of 0.90 and with a low median connectivity indicating the identification of distinct gene modules within the network (**Fig. S2A-E, Table S4**). From the WGCNA, 20 module eigengenes were discovered (MEs) (**Table 2**). Using linear regression, eight MEs were found to be significantly associated with the biofilm state (ME3, ME4, ME5, ME8, ME13, ME16, ME19, ME20) (**Fig. S2G)**.

**Table 2:**
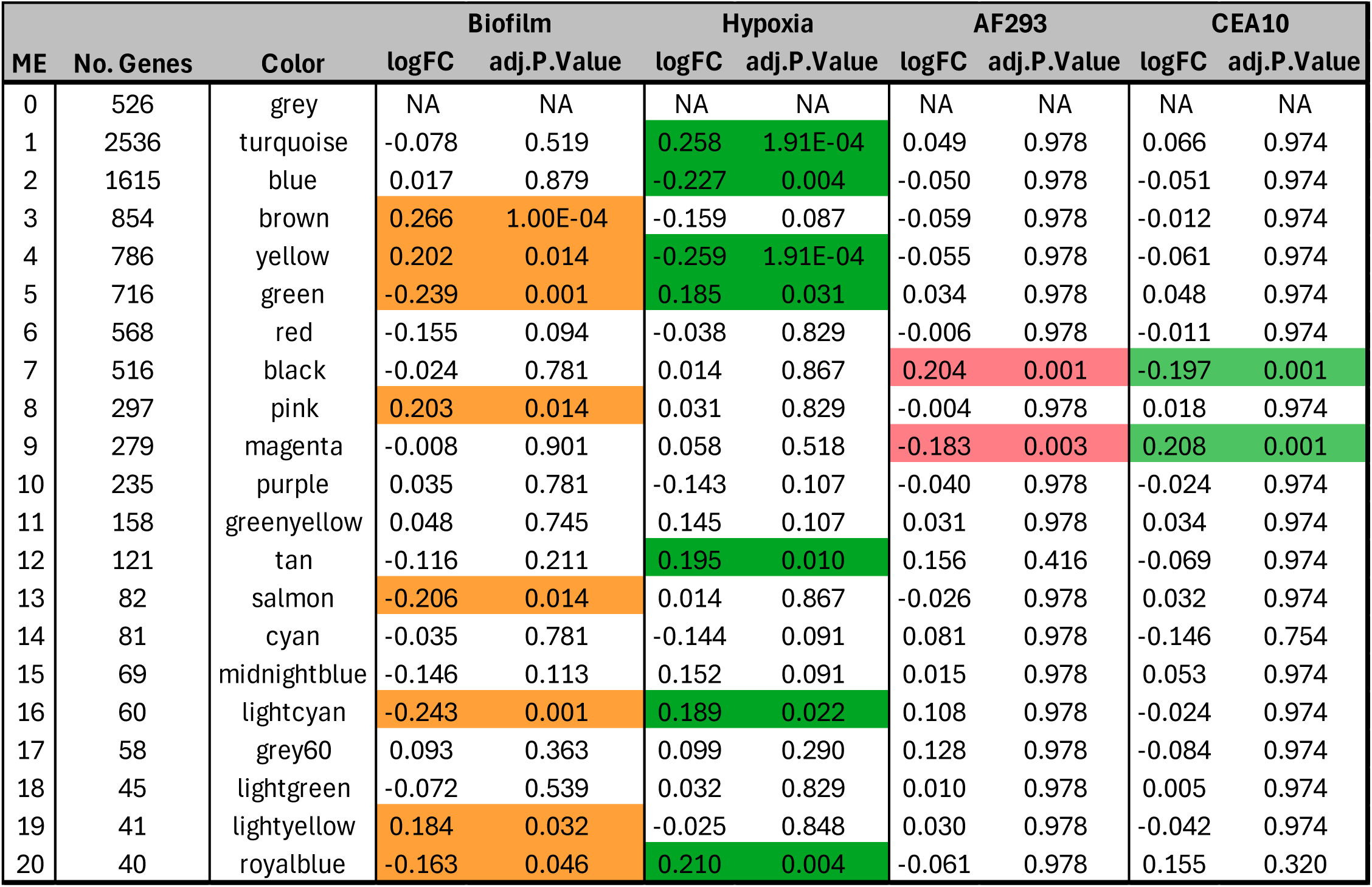
Numbers of genes per module and effect size (logFC) and significance (adjusted p-value) to the biofilm, hypoxia, and strain conditions. MEs that are signficanntly associated (adjusted p-value < 0.05) with a condition are highlighted.

Interestingly, two MEs are significantly associated with *A. fumigatus* strain differences (ME7 and ME9). Previous work has investigated the phenotypic differences between AF293 and CEA10 in the context of colony biofilm growth and pathogenicity (29–31). Functional analysis of ME7 using GO terms found CEA10 and AF293 have alternative regulation of specific transcription factors. Specifically, 52 transcription factor (TF) related genes are differentially regulated between AF293 and CEA10 indicating potential differences in the transcriptional response to the culture conditions examined (**Table S5**). As one example, the TF *stuA* is among the TF genes in ME7. StuA regulates secondary metabolite production and asexual development in filamentous fungi (32–34), and its altered transcript abundance between CEA10 and AF293 may indicate potential differences in secondary metabolite production in the biofilm and potentially explain the differences in asexual development observed between these two common reference strains. Unfortunately, a functional analysis of ME9 does not currently yield new insights into strain specific differences due to the limited annotations of the *A. fumigatus* genome. However, these results potentially highlight the importance of unannotated fungal specific genes under the culture conditions examined.

A surprising finding from the WGCNA analysis is that seven MEs (ME1, ME2, ME4, ME5, ME12, ME16, ME20) are significantly associated with low oxygen conditions. Furthermore, planktonic low oxygen and biofilm shared MEs (ME4, ME5, ME16) contain genes with anti-correlated expression profiles highlighting a biofilm specific hypoxia response (**Fig. S2G**). For example, genes within ME4 have increased abundance in the biofilm compared to the low oxygen planktonic condition with an enrichment for ribosome protein encoding genes. Among these ribosome genes is the ortholog of the *Saccharomyces cerevisiae* RPP0 ribosome protein gene (Afu1g05080) that has a biofilm specific expression pattern with a CPM of 1363 (SD = 252.6) in the biofilm and 210.6 (SD = 154.9) in the planktonic condition.

Among the eight biofilm-specific MEs four contain genes with general increased transcript abundances (ME3, ME4, ME8, ME19) and four contain genes with generally decreased transcript abundances (ME5, ME13, ME16, ME20) (**Fig. 2A**, **Table S4).** Gene Ontology (GO), FunCat, and KEGG enrichment analysis of the MEs with increased mRNA abundance depict a coordinated biofilm response to increase membrane trafficking and secretion (ME3), enrichment for metabolism of lipids (ME3 + ME8), oxidative stress tolerance and redox balancing (ME8 and ME19), RNA splicing and ribosome biogenesis (ME4), and N-glycosylation (ME4) (**Fig. 2A, Table S6**). MEs with reduced transcript abundances in biofilms (ME5, ME13, ME16, ME20) suggest a reduction of polarized hyphal growth with an enrichment for cell cycle and division genes (ME5), reduced mitochondrial activity (ME13), and a reduction of branched chain amino acid and some secondary metabolic pathways (ME16, ME20) (**Fig. 2B, S3A, Table S7**).

**Figure 2:**
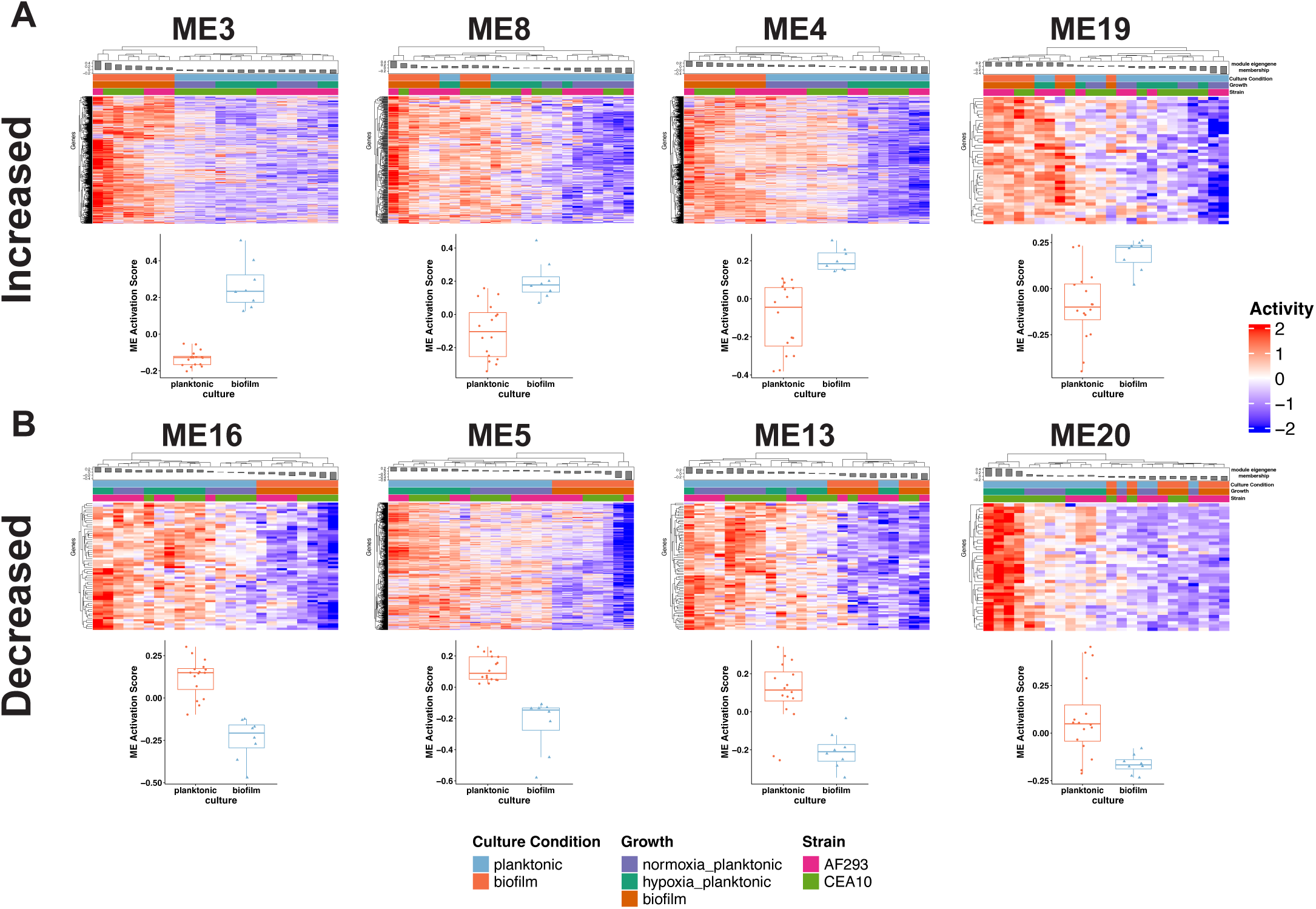
WGCNA reveals eight biofilm specific gene modules. Heatmaps showing transcript abundance levels of genes within the **A**) increased MEs 3, 8, 4, 19 and **B**) decreased MEs 16, 5, 13, 20 with box plots of ME activations score shown below heatmaps. Scaled CPM values are shown in heatmaps.

Together the biofilm MEs portray a biofilm cellular state not focused on polarized growth, cell division and proliferation but rather focused on plasma membrane and cell wall remodeling (increased secretion, lipid metabolism, and transcription), metabolic rewiring, and stress tolerance. Taken together this leads to the hypothesis that cells within the biofilm are forced to adapt to a self-induced stress environment to form and maintain a biofilm. From this we were interested in specific genes that are driving these biofilm adaptations as they may also be involved in generation of emergent biofilm properties such as antifungal drug resistance.

### The highly differentially expressed biofilm specific gene Afu4g06290 is a putative ceramide synthase

Further investigation of genes within the MEs identified the gene Afu4g06290, a putative ceramide synthase, having strong membership within ME3 and standing out as the gene with the strongest gene significance within the dataset (**Fig. 1B, S3A**). Afu4g06290 CPM values are specifically associated with the biofilm state with an average of 15688.08 ± 6058.17 for biofilms and 22.39 ± 7.95 for planktonic cultures (both normoxia and hypoxia conditions) (**Fig. 3A**). In the network analysis Afu4g06290 has a total adjacency score of 73.39 and a degree of 8 using a TOM threshold of 95^th^ percentile of ME3 indicating this gene is both highly significant and also highly centrally located in the ME3 network (**Fig. S3B**). Afu4g06290 has an amino acid sequence similar to the ceramide synthases BarA (AN4332) in *Aspergillus nidulans* (68.62 % identify with 100% coverage) and Bar1 (FGRAMPH1 01 G27129) in *Fusarium graminearum* (44.7% identity with 78% coverage) (35, 36). A phylogeny using ceramide synthase amino acid sequences from a variety of fungi as well as the six human ceramide synthase homologs reveals the *A. fumigatus* Afu4g06290 protein clusters closest with *A. nidulans* BarA and *F. graminearum* Bar1 that produce C18 ceramides (35, 36) (**Fig. S4A**). Interestingly the *Schizosaccharomyces pombe* Lag1 and *Candida albicans* Lac1 are also found in the same cluster rather than with the canonical Lac/Lag proteins ceramide synthase proteins. Additionally, we find that the human ceramide synthase Cers1 protein clusters closer to the fungal ceramide synthases than the remaining human ceramide synthases. Protein folding predictions using the AlphaFold3 server for Afu4g06290, AnBarA, FgBar1 and human Cers1 finds all four structures to be quite similar to Afu4g06290 with ChimeraX matchmaker alignment scores of 1736.4, 1218.7, and 557.8 respectively (**Fig. S4B**) (37–40). Taken together Afu4g06290 is an ortholog of *A. nidulans BarA* and *F. graminearum* Bar1 and will be referred to as BarA (*barA*) hereafter. Previous work showed that loss of *AnbarA* or *Fgbar1* resulted in disruption of polarized growth and growth defect with altered colony biofilm morphology (35, 36). However, the role of *barA* homologs in submerged biofilm models is undefined and its expression profile supported the hypothesis that it plays an important role in the *A. fumigatus* biofilm.

**Figure 3:**
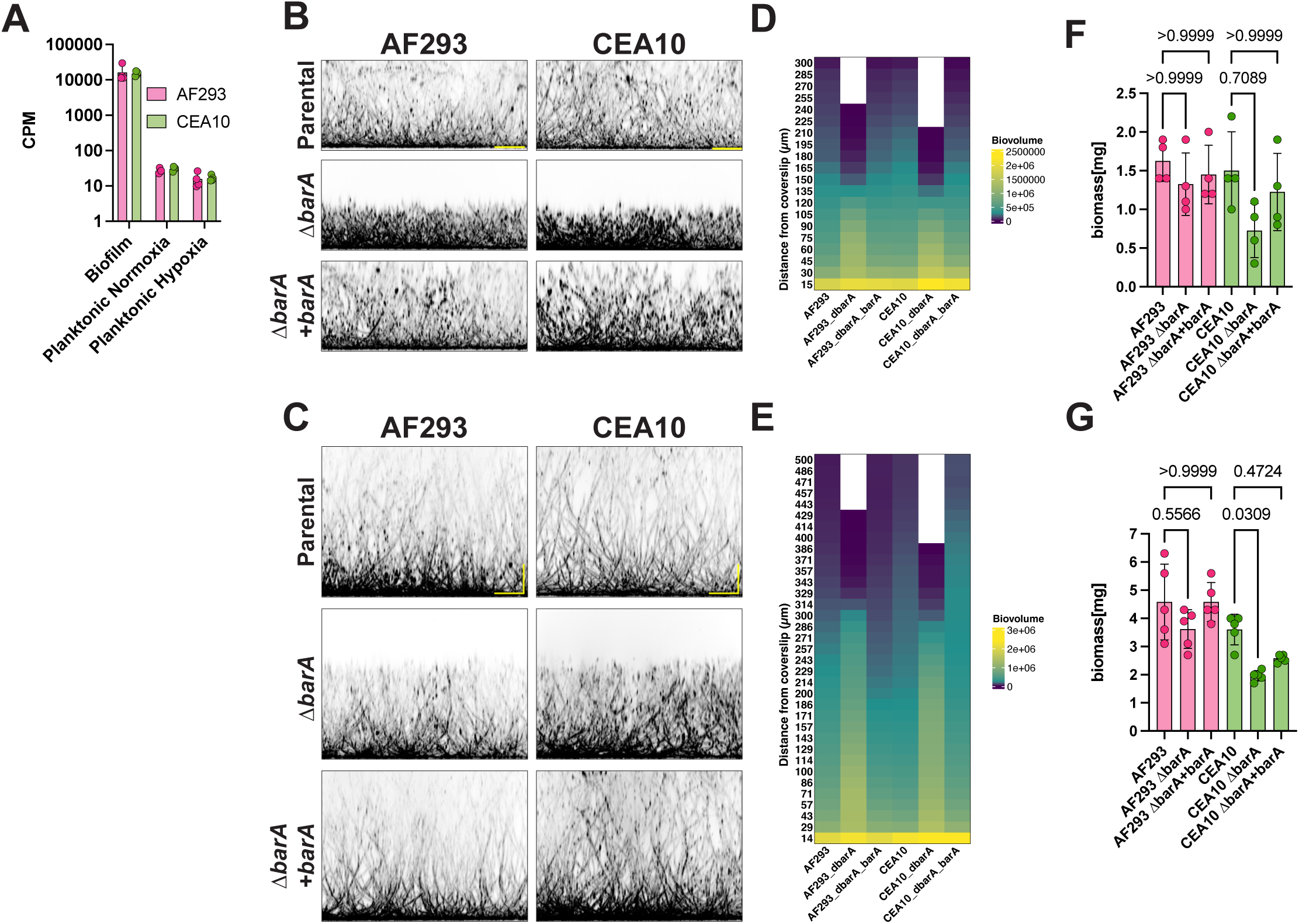
*barA* is required for normal biofilm morphology and oxygen gradients. **A)** CPM values of barA. **B-C**) Microscopy images of calcofluor white stained biofilms were acquired at 18-and 24-hr timepoints and representative XZ maximum projection images of biofilms are shown. Scale bars are 100 µm. **D-E**) Quantification of biofilm biovolume along the height of the biofilm images from n = 3 biological replicates. **F-G**) Total biofilm biomass was measured. Points indicate biological replicates and statistics are a one-way ANOVA with Tukey’s multiple comparison and n = 4-5 biological replicates. Statistical test between wildtype and mutant or reconstituted strain is shown. Statistics are a one-way ANOVA with a Tukey’s multiple comparison test.

### *Aspergillus fumigatus barA* is involved in biofilm formation

With the biofilm specific expression profile, we first hypothesized that BarA is important for biofilm development. We generated *barA* null mutants (Δ*barA)* through gene replacement in both the AF293 and CEA10 reference strain backgrounds. Viable mutants were obtained in both AF293 and CEA10 backgrounds and reconstituted strains were generated by inserting *barA* together with regulatory regions at the aft4 safe-haven locus (41). As reported in *F. graminearum* and *A. nidulans,* the loss of *barA* results in an altered colony biofilm morphology on agar with a reduction in radial growth and a raised rugose colony in AF293 and a raised fluffy colony with irregular edges in CEA10 (**Fig. S4C**). In both strain backgrounds there is a decrease in conidiation in the absence of *barA* (**Fig. S4C**) (35, 36).

Imaging of calcofluor white stained submerged culture biofilms at two timepoints revealed that the loss of *barA* results in a stunted biofilm morphology (**Fig 3B-C**). Biovolume quantification of the biofilms shows that although Δ*barA* biofilms are stunted they are dense as indicated by the increase in biovolume within the central regions of the biofilm compared to the wildtype and reconstituted strains (**Fig 3D-E**). Correspondingly, directly quantifying biofilm biomass at 18- and 24-hours revealed a strain specific difference with the Δ*barA* strains (**Fig 3F-G**). In the AF293 background there is little impact to the overall biomass despite the stunted morphology of Δ*barA*, however, in the CEA10 background, there is a significant reduction in overall Δ*barA* biofilm biomass at the 24-hr timepoint. The reason behind these strain specific differences with loss of *barA* remain to be investigated.

### BarA is involved in biofilm integrity *in vivo*

Given the altered biofilm morphology in the absence of *barA*, we were interested in the impact of BarA on pathogenicity and virulence. We used a murine corticosteroid model of invasive pulmonary aspergillosis (IPA) to investigate the ability of the Δ*barA* mutant to grow within the lung and cause disease. Surprisingly, no significant difference in fungal burden as measured by quantitation of 18S rDNA was observed in the absence of *barA* (**Fig. 4A**). Additionally, histology samples were acquired at the same timepoint as the fungal burden. Interestingly in the hematoxylin and eosin (H&E) staining we found that Δ*barA* lesions stained more readily with eosin (pink color) compared to the wildtype and reconstituted groups (**Fig. 4B**). In addition, we observed that the eosin-stained hyphae were highly vacuolated indicating stressed and potentially dead or dying hyphae. We hypothesized that the Δ*barA* strain could be dying from the center of the lesions in the infection environment. A prediction of this hypothesis is that loss of *barA* would impact disease progression and virulence. However, the Δ*barA* mutant was surprisingly observed to cause the same murine mortality as the wildtype and reconstituted strains (**Fig. S5**). From this data we conclude that BarA produced ceramides likely contribute to the maintenance and viability of a biofilm in the cis-generated hypoxic regions, however, this does not have an impact on host mortality in the setting of an inflammatory host environment and/or acute model of invasive disease, further highlighting the role of the complex infection environment on host mortality in fungal infection models.

**Figure 4:**
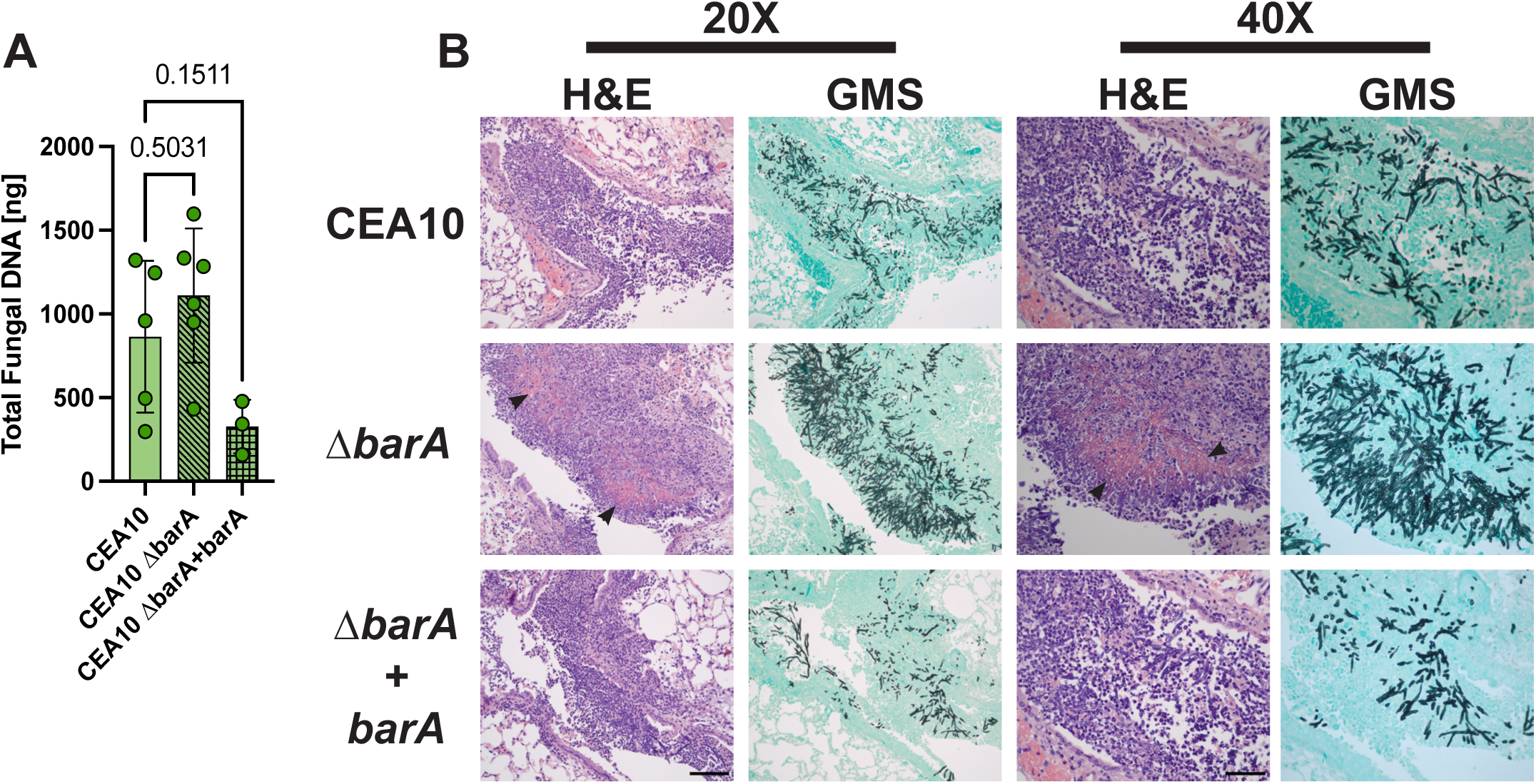
BarA does not impact virulence but does appear to have reduced host fitness. Mice were treated with the triamcinolone on day −1 at 40 mg/kg. On day 0 mice were inoculated with 2×10^6^ conidia per mouse. Mice were euthanized at 72 hours post inoculation and lungs harvested for quantification of fungal burden and histology. **A**) Fungal burden was quantified by qRT-PCR and no statistical difference was observed between the groups. n = 3-6 animals per group. **B**) Lung tissue was prepared for histology and stained with H&E and GMS. Representative images of lung tissue slices at 20X and 40X magnification are shown. Black arrows indicate regions of vacuolation and eosin staining of the fungus. Scale bars are 100 µm for 20x and 50 µm for 40X magnification.

### BarA produces a subset of ceramides

We next sought to understand *barA’s* role in the *A. fumigatus* biofilm. Given the annotation of the *barA* gene product as ceramide synthase, we hypothesized that BarA produces ceramide lipids. To test this, we utilized untargeted lipidomics on AF293 and AF293 Δ*barA* biofilms at 18-hr and 24-hr timepoints. We utilized positive mode LC-MS/MS on lipid extracts generated from AF293 and AF293 Δ*barA* biofilms. We utilized the positive mode to capture the abundance of sphingolipid and ceramide species of lipids given the predicted function of BarA. The technical quality in terms of variability (%RSD) was within expected bounds for the median feature height, and numbers of species features detected. A pooled study sample was generated from all provided samples for quality control purposes, and it was spiked with a mix of isotopically labeled lipids. The intensities and retention times for those standard compounds were observed to generally be consistent across the run indicating good technical data quality. Lipid annotations were performed based on a 3 and 5 ppm search window for MS1 and MS2 feature matching. Annotated features with less than two matched MS2 spectral peaks or a cosine similarity score of less than 0.5 were discarded. This annotation allowed for the annotation of 338 features to lipids in positive mode.

A principal component analysis revealed the first principal component (PC) (32.9% of the data variance) was explained by Δ*barA* versus wildtype (**Fig. S6A**). The second PC showed slight separation based on timepoint mostly observed in the Δ*barA* strain, indicating this PC potentially is explained by some noise in the data. Unsupervised hierarchical clustering of samples and lipids analyzed confirmed that the strain genotype was the main determinate for sample clustering (**Fig. S6B**). K-means clustering of the lipids using five clusters helped to visualize the differences in overall lipid abundance. From this a distinct cluster of lipids containing species of ceramides and glycoceramides was revealed (indicated by the red bracket in the figure) as driving the difference between wildtype and Δ*barA*. A closer inspection of this subset of lipids revealed that *ΔbarA* strain does not produce several C18 ceramides and glycoceramides (**Fig 5A**). Together this analysis supports the conclusion that BarA is a ceramide synthase producing C18 ceramides and C18 glycoceramides in *A. fumigatus* biofilms.

**Figure 5:**
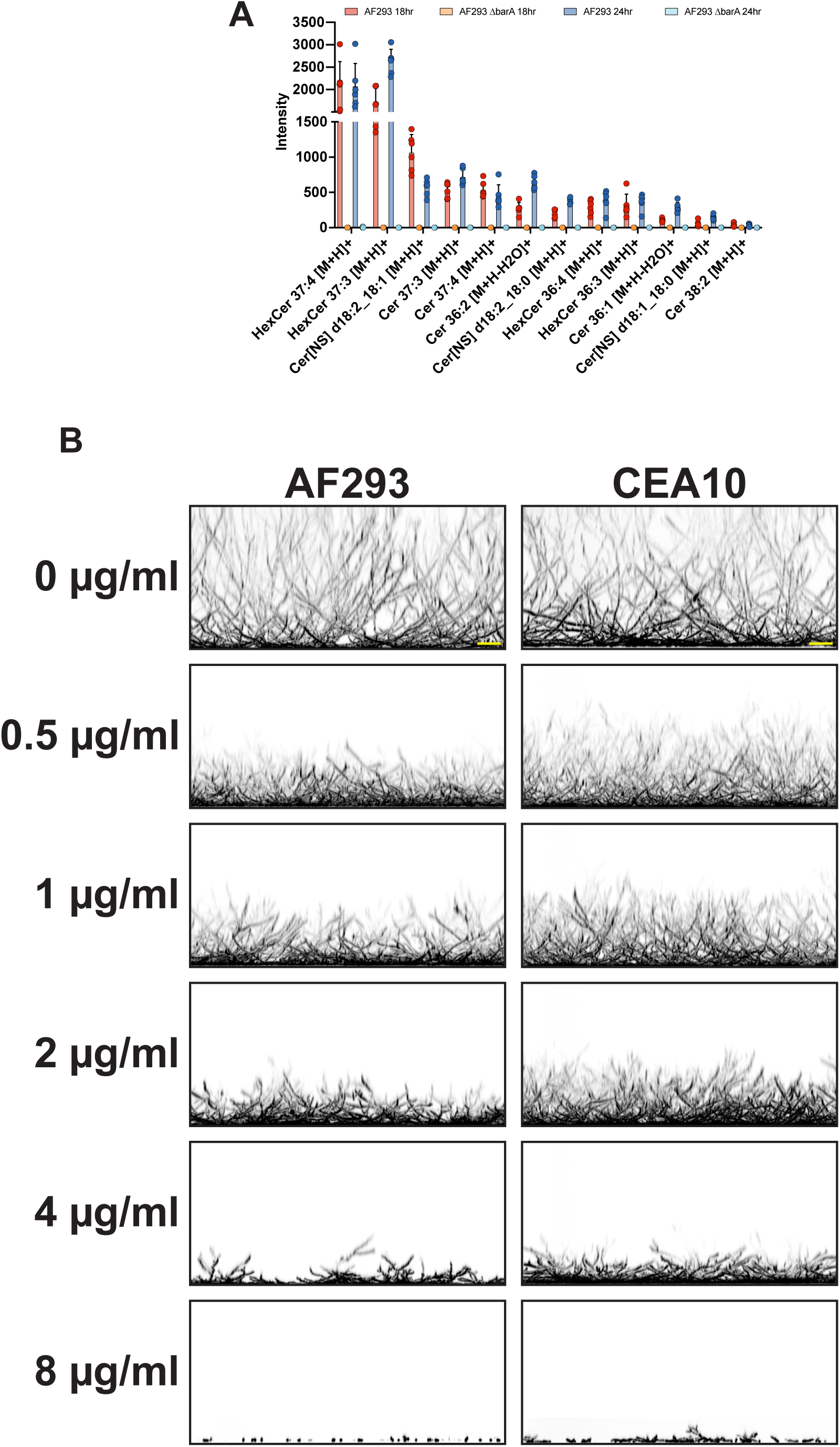
Sphingolipids are important for biofilm formation. **A**) LC-MS/MS was used to profile the lipid content of the Δ*barA* mutant compared to wildtype. A bar plot of a subset of intensity (abundance) values of ceramides identified as being differentially abundant in the heatmap in the Δ*barA* samples from the untargeted lipidomics analysis. The mutant does not produce several ceramides and glycoceramides species. Statistics are a one-way ANOVA with a Tukey’s multiple comparison test. Data points represent biological replicates (n = 6). **B)** Biofilms of AF293 and CEA10 expressing GFP in the cytoplasm were grown for 18 hours in the presence of myriocin at indicated concentrations. Biofilms were imaged using the GFP fluorescence. Images are representative max projected XZ resliced images.

Our findings with the Δ*barA* mutant biofilm morphology and subsequent lipid analysis suggest that sphingolipids and ceramides are important lipid components of a fully functional biofilm. To further test this hypothesis, we treated wild-type *A. fumigatus* biofilms with the serine palmitoyltransferase inhibitor myriocin that inhibits sphingolipid biosynthesis and observed altered biofilm morphology reminiscent of the Δ*barA* biofilm (**Fig. 5B**) (42). Interestingly, myriocin mediated alterations in biofilm morphology are dose responsive indicating that levels of sphingolipids, specifically ceramides, are important for mature biofilm formation. From these data we conclude that BarA produced ceramides are critical for cell adaptation and fitness during dynamic changes in biofilm microenvironmental conditions.

### BarA produced ceramides are important for biofilm antifungal susceptibility

As we observed that the loss of *barA* results in a loss of C18 ceramides and glucosylceramides and impacts biofilm morphology we sought to define the impact of *ΔbarA* on susceptibility to the antifungal drugs that interfere with ergosterol biosynthesis (voriconazole) or target ergosterol itself (amphotericin B). Interestingly, in 18-hr biofilms Δ*barA* mutants have a substantial and significant increase in susceptibility to voriconazole with a ∼50% susceptibility increase in AF293 and ∼80% susceptibility increase in CEA10 compared to the respective wildtype strains as measured by reduction of the metabolic activity dye, XTT (**Fig. 6A, C**). At the 24-hr timepoint there is a slight yet significant increase in susceptibility of Δ*barA* mutants to voriconazole with a ∼22% susceptibility increase in AF293 and a ∼17% increase in CEA10 (**Fig. 6B, D**). The decrease in susceptibility at 24 hours compared to 18 hours in the absence of *barA* suggests that ill-defined dynamic changes in the biofilm cells lacking *barA* partially complement loss of *barA* function.

**Figure 6:**
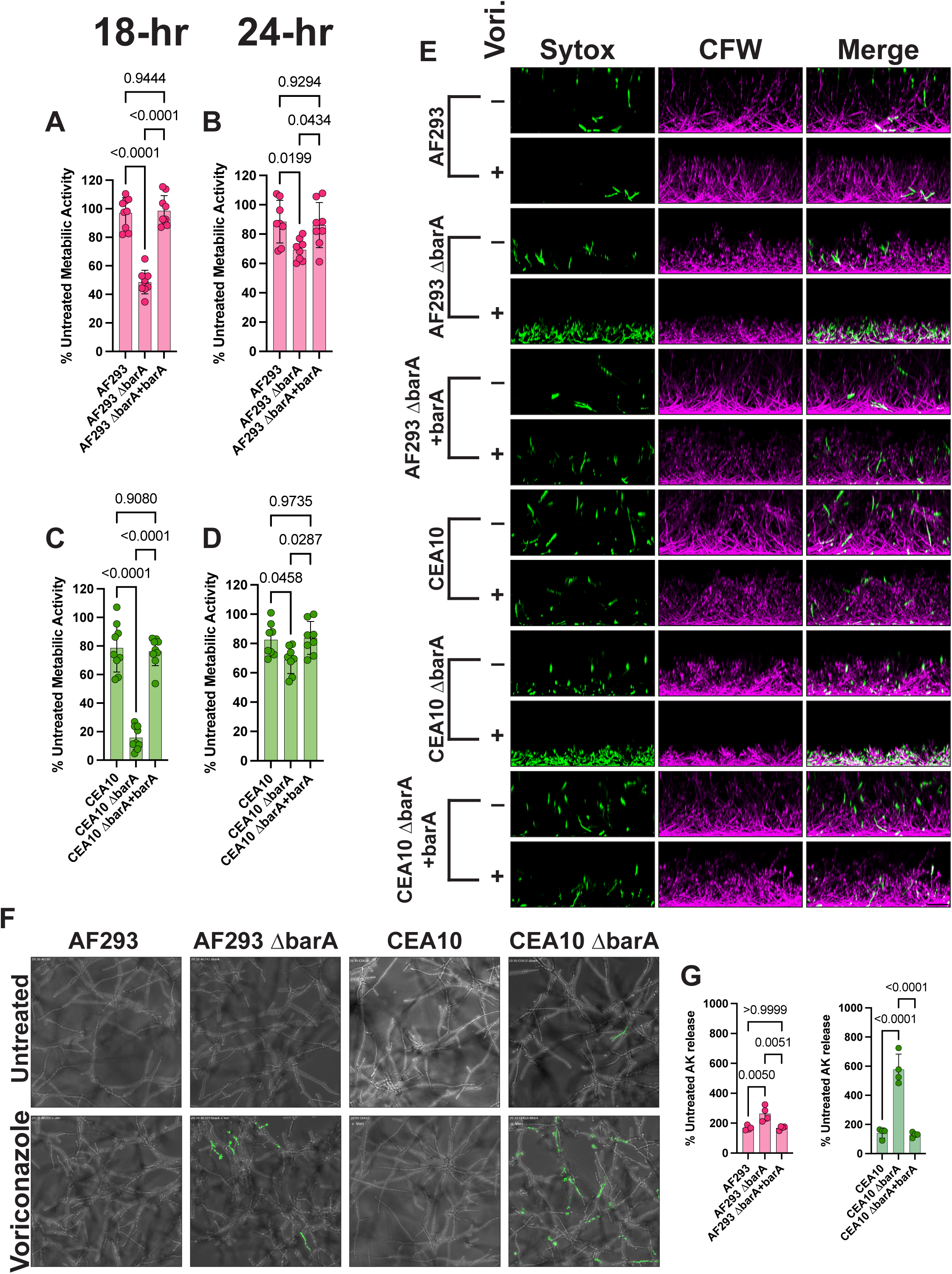
BarA is involved in azole susceptibility. **A-D**) Biofilms at 18- and 24-hours were treated with 1 µg/ml of voriconazole for 3 hours. Metabolic activity was assessed with XTT conversion as a readout of biofilm damage. **E**) Biofilms (18-hour) of indicated strains were treated with 1 µg/ml voriconazole or left untreated for 3 hours. Treated biofilms were stained with calcofluor white (CFW) and Sytox green and imaged to acquire full height of biofilms. Representative images shown are maximum projections of ZX resliced images. **F**). **G**) Quantification of AK release into the media after 3-hour voriconazole treatment of 18-hour biofilms. Statistics are a one-way ANOVA with Tukey’s multiple comparison and n = 6-8 biological replicates for XTT and 3 biological replicates for the AK data.

The impact of voriconazole on Δ*barA* biofilms was explored further through imaging of 18-hour biofilms treated for three hours with voriconazole and subsequently stained with Sytox green (a marker of compromised membranes and dead cells) and calcofluor white (counterstain for all biomass). Consistent with the decrease in XTT reduction, Δ*barA* biofilms have significantly more potentially dead hyphae as indicated by increased Sytox green staining (**Fig. 6E**). With the same voriconazole treatment, the wildtype and reconstituted strains do not show an increase in Sytox green signal. These data suggest that Δ*barA* hyphae have compromised membranes upon azole treatment and that the absence of BarA derived ceramides shifts voriconazole from a fungistatic drug to a fungicidal drug.

To further examine the effect of *barA* loss on voriconazole biofilm susceptibility, we needed to alter our approach as our full height biofilm images do not provide sufficient resolution to assess the reason for the observed Sytox staining due to low magnification and observing a single timepoint of three hours. To address this technical limitation, the biofilms were subsequently imaged at a higher magnification using timelapse microscopy before and during antifungal treatment. In these experiments, only the bottom 30 µm of the biofilm were imaged. Again, Sytox green was utilized to indicate cells with compromised membranes. In these time lapse movies beginning approximately two hours after addition of drug, Sytox staining is observed and continues to increase up to three hours in both Δ*barA* strain backgrounds (**Movies S1,S2**). End point images are represented in **Fig. 6F**. The observed Sytox staining is due to cell lysis events as cytoplasmic contents are observed being released into the milieu. Consequently, voriconazole treatment at 18-hours of biofilm development is leading to catastrophic cell lysis in the absence of *barA*. Importantly in these experiments there was little Sytox staining observed in the wildtype treated and untreated samples consistent with what is observed in the full height imaging in **Fig. 6E**.

Additionally, *ΔbarA* mutants biofilms were tested for susceptibility to the polyene antifungal drug amphotericin B. In contrast to voriconazole, at both timepoints and in both strain backgrounds loss of *barA* confers a significant reduction in susceptibility to amphotericin B (**Fig. 7A**). In the AF293 background Δ*barA* has a ∼220% susceptibility decrease and ∼65% susceptibility decrease in CEA10 at the 18-hr timepoint. At the 24-hr timepoint there is a ∼119% susceptibility decrease in AF293 and ∼72% decrease in CEA10. Interestingly, the impact of the Δ*barA* mutant on susceptibility is stronger at 24-hr compared to the 18-hr timepoint. To more directly assess the damage to the biofilm we also quantified the amount of adenylate kinase (AK) released into the media after treatment with voriconazole and amphotericin B at the 18-hour timepoint. In agreement with our XTT data we found more AK release in Δ*barA* biofilms treated with voriconazole (**Fig. 6G**) and less AK release in Δ*barA* biofilms treated with amphotericin B (**Fig. 7B**). Taken together these data indicate that BarA is involved in ergosterol dynamics in biofilm cells and is an important mediator of biofilm antifungal drug susceptibility.

**Figure 7:**
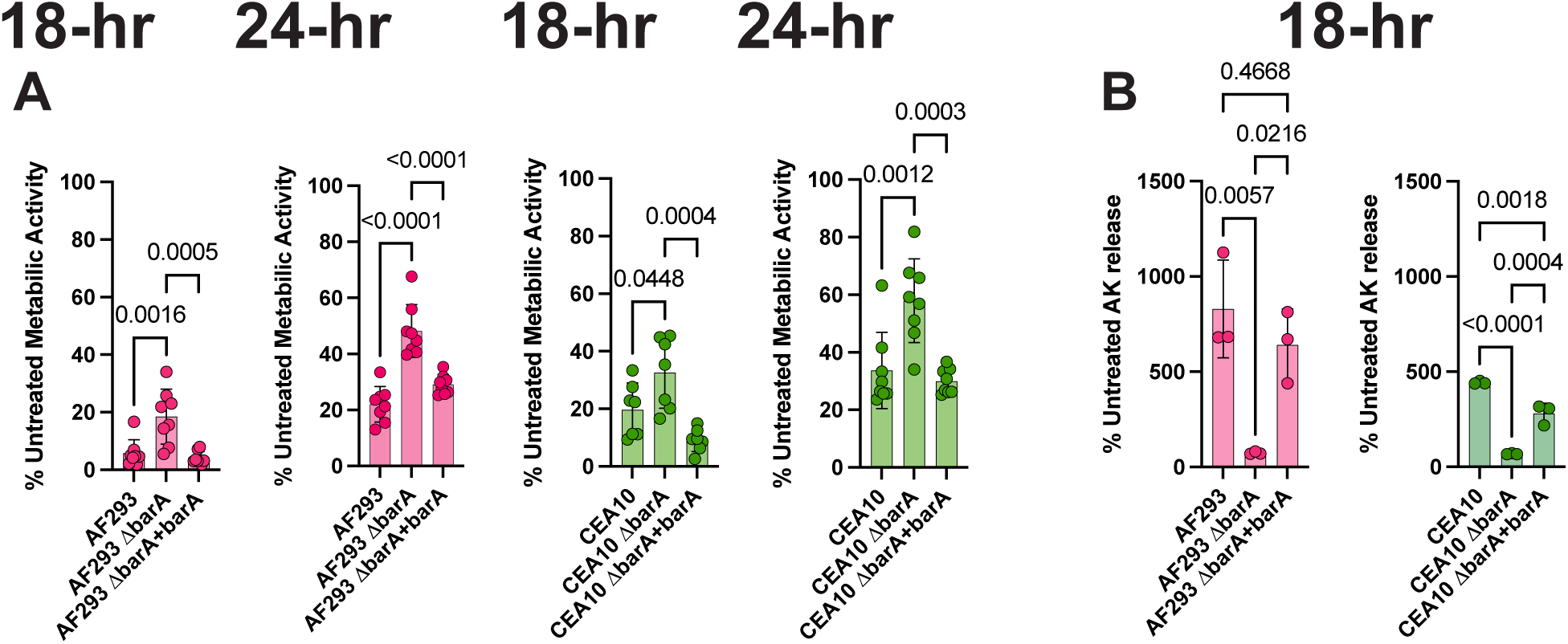
Δ*barA* biofilms have a reduced susceptibility to amphotericin B. **A**) Biofilms were grown to indicated time and treated with 1 µg/ml amphotericin B for three hours. Metabolic activity was assessed with XTT conversion. **B**) Biofilms were grown to indicated time and treated with 1 µg/ml amphotericin B for three hours and AK release was assessed. Statistics are a one-way ANOVA with Tukey’s multiple comparison and n = 7-8 biological replicates for XTT and 3 biological replicates for the AK data.

### BarA derived ceramides are important for ergosterol localization

The decrease in susceptibility to the ergosterol targeting drug amphotericin B observed in the Δ*barA* mutants suggests that these strains have less ergosterol localized to the plasma membrane. Additionally, the increased sensitivity to voriconazole suggests that Δ*barA* strains have increased sensitivity to alterations in available ergosterol. Ceramides, specifically C18 derived glucosylceramides, have been implicated in the proper localization of ergosterol to the plasma membrane (35, 36). These lipids have been proposed to bind with ergosterol and form detergent-resistant lipid rafts (43). The addition of the glucose moiety to the C18 ceramide allows for these ergosterol rich rafts to expand into large domains important for polarized growth and stress tolerance (43).

To determine if plasma membrane ergosterol is reduced in Δ*barA* strains, we utilized filipin to examine the early stages of biofilm growth initiation (germlings). As there is an enrichment of ergosterol in the growing tip we decided to use germlings to investigate the level of ergosterol in the tip of the Δ*barA* mutants compared to the wildtype strains. Staining was performed on live cells as previously reported (35). In support of our hypothesis and previously published data in other fungi, we observed that the loss of *barA* leads to a decrease in filipin localization at the growing tip in both strain backgrounds (**Fig. S7**). Importantly observing the population of germlings, there are fewer cells with increased filipin staining in the tip in the mutant strains while the wildtype strain had very strong tip staining in the majority of germlings indicating loss of tip localized ergosterol in the Δ*barA* mutants. We further quantified the intensity of tip localized filipin stain from our images. From this quantification we found a significant reduction in the intensity of tip localized filipin in the Δ*barA* in both strain backgrounds with a 27.7% reduction in AF293 and a 34.8% reduction in CEA10. This data strongly indicates the BarA produced ceramides play a role in ergosterol dynamics and therefore antifungal susceptibility within the *A. fumigatus* biofilm.

## Discussion

Historically the transcriptional profiles of *A. fumigatus* have been in the context of homogenous culture conditions. This has provided a wealth of information regarding the biology of this important pathogen but has fallen short on describing the transcriptional space of the host relevant growth mode of a biofilm. Previous work investigated the emergent properties of the *A. fumigatus* biofilm; however, questions remained in how the biofilm differs transcriptionally from other growth states (9). Here we have directly addressed this question by comparing the transcriptional state of homogenous planktonic batch cultures to that of the biofilm. Due to previous work observing that the biofilm has self-generated low oxygen gradients we included a planktonic group that was exposed to low oxygen. The purpose of this group was to allow us to study the biofilm specific transcriptional responses away from the general low oxygen responses that have previously studied (23–26). From this analysis we observed that the biofilm occupies a unique transcriptional state from cultures grown in atmospheric oxygen levels and from cultures exposed to a low oxygen environment. Importantly, we have seen that despite strain specific differences between the two laboratory strains used in this study there is a robust biofilm specific response. Previous work has exemplified the phenotypic and virulence differences between the two *A. fumigatus* reference strains, AF293 and CEA10, with reference genomes (29, 30, 44). Here we found that the two strains have differences in the abundance of a significant number of transcription factors. Furthermore, previous work has demonstrated differences in genomic content in the form of accessory genes between *A. fumigatus* (45–47). These differences likely have significant contributions to the observed strain specific phenotypes (29, 30, 44). In the context of the current study the robustness of the biofilm specific expression patterns is increased given the heterogeneity between AF293 and CEA10. While we cannot say whether these observed differences contribute to the differences in virulence of AF293 and CEA10, these data provide a foundation to continue exploration of mechanisms behind strain specific pathogenicity and virulence.

A major goal of this study was to identify novel genes important for biofilm fitness and antifungal drug responses. To help identify biofilm specific genes, we built on the initial differential expression analysis to improve granularity through a WGCNA with the hypothesis that we would identify modules of genes with biofilm specific gene expression patterns. Functional ontology analysis of these biofilm gene modules paints a picture of a gene expression pattern of metabolic rewiring, altered cell cycle, and cell division in the biofilm compared to the planktonic state. The inclusion of the low oxygen planktonic batch culture condition allowed us to uncover that the biofilm culture model is responding to the low oxygen gradients that develop within the biofilm in a different manner than planktonic cells exposed to low oxygen. It is likely the biofilm response represents gradual adaptations to a dynamic nutrient and osmotic environment as low oxygen zones develop in contrast to the shock of a rapid switch from 21% to 0.2% O_2_.

We chose to focus on ME3 due to the finding of a highly connected gene (*barA)* within this network. However, the eight biofilm specific modules provide a further understanding of the biofilm suggesting that metabolic shifts such as an increase in fermentation leading to acetyl-coA and a reduced focus on polarized growth and cell division are important and await further investigation. An additional theme of the genes found in these eight MEs is a general stress response indicating the biofilm develops self-imposing stress that requires adaptation for persistence. Exemplifying this point further, the gene *ishA* (Afu3g10480) is found in ME3. In the fission yeast *Schizosaccharomyces pombe,* ish1 is involved in survival of stationary phase, glucose starved cells (35). It is likely that the cells in the most interior regions of the biofilm experience stationary phase like conditions with local nutrient depravation in addition to significantly reduced oxygen availability. Further studies will focus on the biology of these cells and understanding the role they play in the emergent properties of a biofilm. One potential avenue is spatial transcriptional mapping of the biofilm using marker genes identified in biofilm specific MEs.

Using our transcriptional analysis, we sought to identify target genes that are important for biofilm form and function. We hypothesized that genes with strong biofilm specific module membership, high network centrality and robust differential expression patterns would be good targets. Using this logic we identified the gene *barA* (Afu4g06290) as having a highly specific and robust increase in transcript abundance in biofilms compared to both planktonic conditions examined and strong module membership to the biofilm specific ME3. BarA significantly contributes to the morphology of the *A fumigatus* biofilm (colony and submerged culture). Interestingly, the overall submerged biofilm biomass produced in the absence of *barA* was only slightly impacted and was strain dependent. These data suggest a component of the strain specific differences observed in our study is related to lipid metabolism and membrane homeostasis.

Accordingly, BarA generates C18 ceramide lipids. Glucosylceramides derived from C18 ceramide have been proposed to aid in the proper localizing of ergosterol in the plasma membrane through formation of expandable lipid rafts containing glucosylceramide and ergosterol (48). Therefore, the likely mechanism of antifungal susceptibility differences in the Δ*barA* strains is from a lack of glucosylceramides that alter ergosterol membrane localization.

Additionally, the alterations in ergosterol localization likely explains the polarity differences that lead to the observed altered biofilm morphology. Without the proper localization of ergosterol, the maintenance of polarized growth is impacted resulting in increased hyphal branching and a shorter, but denser, biofilm.

Another potential and attractive role of C18 ceramides is in potentially regulating cell metabolism or quiescence. In mammalian systems C18 ceramide has been shown to be involved in regulating LC3 mediated mitophagy (49). Therefore, a potential explanation for the dramatic increase in *barA* transcript levels in the biofilm is the microenvironment unfavorable to mitochondrial mediated respiration. C18 ceramides may be needed to inactivate mitochondria specifically in basal depths of the biofilm, though this remains to be experimentally tested. In support of this hypothesis, work in *Aspergillus nidulans* found that basal level cells in the biofilm show dispersal of microtubules in later stages of biofilm development when oxygen tensions in the biofilm are low (50, 51). The dispersal of microtubules in mature biofilms was dependent on SrbA and supports the hypothesis that basal level cells are entering a quiescent-like state (50, 51).

A surprising finding was that BarA is dispensable for virulence in a corticosteroid murine model of invasive pulmonary aspergillosis. However, in further support of the submerged biofilm model being predictive of *in vivo* growth, the Δ*barA* sites of infection in the murine lung have a similar increase in hyphal density as was observed in the microscopy of the *in vitro* biofilms. A clear difference in the infection site with Δ*barA* compared to the wild-type and reconstituted strain is increased eosin staining within the dense fungal lesions. Moreover, hyphae within these eosin positive lesions contained large numbers of what appears to be vacuoles. It has previously been reported that 4-day old cultures of the *Fusarium graminearum* ΔBAR1 showed a striking increase in vacuolation and aberrant morphology that was not due to cell death and also did not impact pathogenicity (35). While immune cellularity was not quantified, analysis of the H&E images did not suggest an increase in eosinophils to Δ*barA* lesions. It is possible that the Δ*barA* cell surface or secreted matrix/proteins are altered. Additionally, Δ*barA* could be inducing a different type of host cell death leading to increased eosin staining. As fungal burden is measured by quantitation of 18S rDNA, we cannot rule out viability differences at the site of infection with loss of *barA.* Among the challenges of studying disease progression in murine models of IPA is the acute time course of mortality in these models. It seems likely in a chronic or less acute disease model that loss of *barA* would impact virulence. Thus, while no difference in murine survival was observed with mice challenged with Δ*barA* compared to isogenic controls, we cannot rule out that the cause of mortality is different with Δ*barA* infections. Additionally, future studies are needed to investigate the potential role of *barA* in antifungal resistance in the context of the infection environment.

Taken together we observed that the *A. fumigatus* biofilm transcriptome is distinct from planktonic batch cultures and highlighted by transcripts involved in cell growth and stress responses. Intriguingly, while the two reference strains share many features of this biofilm specific response, significant unexplained differences were observed that are worth investigating in future studies. A conserved response was the large induction of the gene *barA* critical for C18 ceramide lipid biosynthesis. Loss of *barA* impacted the formation of an *A. fumigatus* biofilm and the emergent property of antifungal drug resistance, again with strain specific effects observed. These data suggest that in part the strain specific differences are related to lipid and membrane homeostasis. Finally, our study not only provides insight into the biology of *the A.* fumigatus biofilm structure but sets future studies up to interrogate the biofilm at a more granular level to investigate transcriptional states at a spatial level within the biofilm structure of diverse strains.

## Materials and Methods

### Wild type strains, media and growth conditions

Wildtype strains used in this study were AF293 and CEA10 (also known and FGSC A1163). Conidia were generated a previously described (52). Media for experiments was a synthetic complete, nitrogen, glucose media (SCN) designed for this study. SCN contains 1% glucose, 2 g/l SC mix (Sunrise Science), 1.46 g/l glutamine, 0.51 g/l NaNO3, 0.52 g/l KCl, 0.52 g/l MgSO4 7H2O, 1.52 g/l KH2PO4 monobasic, 2.2 mg/l ZnSO4 7H2O, 1.1 mg/l H3BO3, 0.5 mg/l MnCl2 4H2O, 0.5 mg/l FeSO4 7H2O, 0.16 mg/l CoCl2 5H2O, 0.16 mg/l CuSO4 5H2O, 0.11 mg/l Na2MoO4 2H2O, 5 mg/l Na4EDTA, 1% glucose; pH 6.5. Cultures were grown at 37 °C + 5% CO_2_.

### Sample preparation for RNAseq

For biofilms static cultures were seeded in SCN with 10^5^ spores per mL in three wells of a six well culture dish using 2ml per well, while planktonic cultures were grown in 100 ml of SCN at 10^5^ spores per mL. Biofilm and normoxia planktonic shaking flask cultures were grown for 18 hours at 37 °C + 5% CO_2_ prior to tissue collection, while planktonic shaking flask hypoxia cultures were grown for 18-hours and switched into 0.2% O_2_ + 5% CO_2_ for an additional 30-mintues shaking. For biofilms biomass was collected by removing media and replacing with 1 mL with TRI Reagent (Invitrogen) and collecting tissue on ice. Shaking biomass was collected via filtration through miracloth and immediate placed in TRI Reagent on ice. Biomass spun and TRI reagent was replaced with fresh 200 µl TRI reagent and bead beat with 2.3 mm silica beads. The aqueous phase of TRI reagent was obtained by following manufacturer’s protocol, precipitated with 70% ethanol and loaded on a RNeasy column (Qiagen). RNA was collected from columns following manufacture’s protocol. RNA samples were DNAse treated with Turbo DNA-free kit (Invitrogen).

### RNA-sequencing

RNA for RNA-seq was quantified by qubit (Thermo Fisher Scientific) and integrity measured on a fragment analyzer (Agilent). Samples with RIN ≥ 7 underwent library preparation with the mRNA HyperPrep kit (Kapa Bioscience) using 200ng (biofilm) or 100 ng (planktonic) RNA as input following manufacturer’s instructions. Libraries were pooled for sequencing on a NextSeq500 instrument (Illumina), targeting 10M, single end 75bp reads/sample for biofilm samples and a Nextseq2000 (Illumina) targeting 10M, paired end 50bp reads/sample for planktonic samples.

### RNAseq analysis

Raw read quality was assessed using fastQC v. 0.11.8 (Babraham Bioinformatics group) and multiQC v. 1.10.1 software (53). Reads were trimmed using Cutadapt v. 2.4 software using a quality score cutoff of 20 (54). Alignments were performed using Star alignment software v. 2.7.2b (55) with the *A. fumigatus* AF293 reference genome version FungiDB-52 and general feature format (GFF) file from same version. Counts per gene were compiled using HTseq-count v. 0.11.2 (56). R version 4.4.2 was used for the differential expression and network analysis. For the exploratory analysis the package DESeq2 v. 1.46.0 (57) was used to generate vst normalized values for use in clustering and principal component analysis. The package edgeR v. 4.4.2 (58) was used to normalize read data for use in differential expression analysis using the TMM method. Low abundant transcripts were filtered using the fitlerByExpr function and normalization was performed using the CalcNormFactors with using trimmed mean of the M-values. Limma-voom v. 3.62.2 methodology was used to perform a differential expression analysis using linear model fit with Empirical Bayes moderated t-statistics as previously described (59) while controlling for condition as a random effect. Scaled CPM values are used for heatmap representation using the R scale function which works by subtracting values from the mean CPM for each gene and dividing by the standard deviation for the same gene. Heatmaps were generated using the ComplexHeatmap 2.22.0 package (60). WGCNA v. 1.73 was performed as described (27, 61). For our network we utilized a soft threshold of 20 and a tree cut height of 0.25. kME (module eigengene-based connectivity), which represents the correlation between a genes expression profile and the first principal component of a given module’s gene expression matrix, was used to assess strength of module membership and identify potential ‘*hub’* genes (highly connected genes that are centrally located within a module). kME values range between 0 and 1, with higher values indicating greater module membership and intramodular connectivity. The gene set enrichment analysis was performed using FungiFun3 web based software (https://fungifun3.hki-jena.de/) using defult settings using the *A. fumigatus* AF293 species (62). The input were all genes with a adjusted p-value of less than 0.05 from the differential expression analysis. Functional ontology enrichment analysis for differentially expressed gene and ME gene lists was also performed using FungiFun3 software using defult settings and the *A. fumigatus* AF293 species.

### Protein sequence comparison and structure prediction

Protein sequence comparison was made using the EMBL-EBI dispatcher system with an Clustal Omega multiple sequence. The structure of BarA and related orthologs was modeled using the Alphafold3 server (37). Models were displayed visually and compared using ChimeraX software (40). Matchmaker with standard settings was used to determine the structural similarity of the predicted models in a pairwise fashion.

### Strain Construction

The Δ*barA* mutant was generated by gene replacement with the *ptrA* pyrithiamine resistance dominant selection marker gene. The transformation utilized the CRISPR/cas9 method of transforming protoplasts as previously described (52, 63). Mutants were selected for on osmotically stabilized media containing pyrithiamine (100 µg/l). Loss of *barA* was confirmed by PCR and southern plot using a DIG labeled probe for the *ptrA* selection marker on single spored mutant candidates. The *barA* reconstituted strain (*ΔbarA + barA*) was generated by construction of a plasmid containing the *barA* ORF, 1211 bp upstream, 811 bp downstream of the ORF, and the hphA dominate selection marker gene for hygromycin resistance. The reconstitution cassette was integrated at the *aft4* safe-haven locus (41). Expression levels of the *ΔbarA and ΔbarA+barA* strains were confirmed by reverse transcriptase quantitative PCR using RNA extracted from 18-hour biofilms.

### Microscopy and sample preparation

Images were acquired on a Nikon spinning disk confocal microscope equipped with a Yokogawa CSU-W1 spinning disk head and 20X, 40X, or 60X magnification as indicated with 488nm and 405nm lasers and transmitted light. Images were captured on either a Zyla sCMOS (Andor) or a Prime BSI (Teledyne) detector. Fluorescent strains were imaged directly while calcofluor white stained biofilms were stained with 25 µg/ml of calcofluor white 20 minutes prior to imaging. Sytox Green staining was used with a final concentration of 1 µM. Filipin staining was performed by addition of 25 µg/ml of filipin complex (Sigma-Aldrich) to germling cultures incubating for 3 minutes and acquiring images for 7 minutes. Time lapse images of Sytox staining was performed with Sytox Green (1 µM) in culture from the start of the experiment. For myriocin, biofilms were grown with myriocin (Sigma-Aldrich) for 18 hours and stained with calcofluor white for imaging as above.

### Microscopy analysis

Image processing and representation was performed using Fiji software (64). Analysis of filipin staining was performed by drawing a 10-pixel wide line along the length of the hyphae through the tip and maximum value determined. The maximum value was subtracted from the background nearby. Biofilm biovolume analysis was performed as previously described using BiofilmQ software (8, 9, 65).

### Untargeted LC-MS Lipidomics Instrumental Analysis

Samples were prepared by growing biofilms to indicated times in SCN media. Biofilms were washed 3 times with fresh 75mM ammonium carbonate (pH7.4 with acetic acid). Biomass was collected, centrifuged briefly, supernatant removed and pellet flash frozen in liquid nitrogen. Lipid extraction was performed on same day for all biological replicates by resuspending thawed pellet in 800 µl of extraction buffer (1:1 LC/MS grade isopropanol and methanol), and bead beating 1:30 minutes. Debris was pelleted by centrifugation and supernatant (metabolite extracts) was collected for analysis. Metabolite extracts were analyzed by LC-MS/MS. 2 μL of each sample was injected and separated by UHPLC using a Nexera UHPLC system (DGU-405 degasser unit, LC40DX3 solvent delivery system, SIL-40CX3 auto sampler, CBM-40 system controller, CTO-40C column oven; Shimadzu). Separation was achieved by reverse-phase liquid chromatography using an Acquity UPLC BEH C18 column (1.7 μm, 2.1 mm X 30 mm; 186002349, Waters). Separation was achieved using a 5-minute multi-phase linear gradient of the following buffers: Buffer A) 6:4 acetonitrile: H2O (v/v) 1 mM ammonium acetate, B: 9:1 isopropyl alcohol: acetonitrile (9:1, v/v) with 1 mM ammonium acetate. Samples were ionized using an Optimus Turbo V + Dual TIS ion source (Sciex) and were analyzed using an X500R mass spectrometer (Sciex). Samples were analyzed in positive ionization mode and were acquired using a high-resolution data dependent acquisition method. An exclusion list was developed through two serial top-16 acquisitions on a pooled study standard. In the main run a top 6 method was used.

### Untargeted LC-MS Lipidomics Data Processing

LC-MS/MS data processing and ion annotation was performed according to accepted protocols for mass spectrometry data processing and feature annotation. Briefly, annotation was based on matching of chromatographic retention times, MS1 values from detected features, and the resulting MS2 fragment ions to expected retention time ranges for the indicated lipid classes. Annotation was provided as MS1/MS2 when a cosine similarity score of >0.5 with at least two matched signals was observed. An m/z difference of 5 mDa or 3 ppm was allowed for precursor m/z matched, and fragments were allowed a 10 mDa or 5 ppm m/z difference. For MS1/rt matches, a 3 mDa or 3 ppm m/z tolerance was allowed, and the signal needed to elute within 0.3 min of the expected retention time for that lipid or lipid class. Observed peak heights were normalized to the sum of total annotated feature intensities per sample.

### Biofilm susceptibility studies

Biofilms were grown and treated as previously described (9). Briefly, biofilms for this study were grown and treated in SCN media and 1µg/ml of voriconazole or amphotericin B was used.

### Murine models of pathogenesis

#### Fungal burden and histology

A murine model of invasive aspergillosis was used as previously described (8, 30). Female CD-1 outbreed mice at 20-24g were used (Charles River Laboratory). Mice were immunosuppressed with a single dose of triamcinolone acetonide (Kenalong-10, Bristol-Myer Squibb) at 40 mg kg^-1^ one day prior to inoculation. Inoculation was performed intranasally as previously described with 2×10^6^ conidia per 40 µl of PBS (8, 30). Lungs were harvested at 72 hours post inoculation and prepared for fungal burden and histology as previously described (8, 66). *Survival.* Immune suppressed mice were inoculated intranasally with 10^5^ conidia per 40 µl PBS. Mice were monitored for morbidity and endpoint criteria as previously described (30, 66). Kaplan-Meier curves were generated, and a Mantel-Cox log-rank and Gehan-Breslow-Wilcoxon tests were used to test for significance.

### Statistical Analysis

Statistical analyses for figues 3-7 and S5,6 utilized GraphPad Prism 10 software. Error bars in plots indicate standard deviation around the mean.

## Data availability

The data discussed in this publication have been prepared to depost in NCBI’s Gene Expression Omnibus (67) and will be deposited upon re-opening of the website.

## Code availability

R script for differential expression analysis and WGCNA are available on GitHub (https://github.com/charlespuerner/AFUM_Biofilm_vs_Planktonic).

## ACKNOWLEDGEMENTS

This work was funded by NIH/NIAID grants R01AI130128 (RAC) and R01AI146121 (RAC). Core facility support provided by NIH grant P20-GM113132 to the Dartmouth BioMT COBRE, the Genomics Data Sciences Core at Dartmouth, supported by the National Institutes of Health-funded Center for Quantitative Biology at Dartmouth (COBRE, 5P20GM130454-03) and the Dartmouth Cancer Center core grant (P30CA023108) from the National Cancer Institute. Additional support was provided by the Cystic Fibrosis Foundation Research Development Program (STANTO19R0) and NIH/NIDDK P30-DK117469 (Dartmouth Cystic Fibrosis Research Center). The authors would also like to thank Ann Lavanway in the Dartmouth Microscopy core for assistance with imaging and Dr. Carey Nadell at Dartmouth for training on BiofilmQ image analysis.

---

**Movie S1:** Timelapse images of AF293 and AF293 Δ*barA* biofilms untreated or treated with 1 µg/ml of voriconazole for 3 hours. Wildtype untreated is upper right, wildtype treated is upper left, mutant untreated is bottom left and mutant untreated is bottom right. The timepoint where the treatment is added is indicated in the labeling.

**Movie S2:** Timelapse images of CEA10 and CEA10 Δ*barA* biofilms untreated or treated with 1 µg/ml of voriconazole for 3 hours. Wildtype untreated is upper right, wildtype treated is upper left, mutant untreated is bottom left and mutant untreated is bottom right. The timepoint where the treatment is added is indicated in the labeling.

## References

1. Monds RD, O’Toole GA. 2009. The developmental model of microbial biofilms: ten years of a paradigm up for review. Trends Microbiol 17:73–87.

2. Katharios-Lanwermeyer S, O’Toole GA. 2022. Biofilm Maintenance as an Active Process: Evidence that Biofilms Work Hard to Stay Put. J Bacteriol 204.

3. Sauer K, Stoodley P, Goeres DM, Hall-Stoodley L, Burmølle M, Stewart PS, Bjarnsholt T. 2022. The biofilm life cycle: expanding the conceptual model of biofilm formation. Nat Rev Microbiol 20:608–620.

4. Gondil VS, Subhadra B. 2023. Biofilms and their role on diseases. BMC Microbiol 23:1–3.

5. Ramage G, Rajendran R, Sherry L, Williams C. 2012. Fungal Biofilm Resistance. Int J Microbiol 2012:528521.

6. Morelli KA, Kerkaert JD, Cramer RA. 2021. Aspergillus fumigatus biofilms: Toward understanding how growth as a multicellular network increases antifungal resistance and disease progression. PLoS Pathog 17:e1009794.

7. Desai J V., Mitchell AP, Andes DR. 2014. Fungal Biofilms, Drug Resistance, and Recurrent Infection. Cold Spring Harb Perspect Med 4:a019729.

8. Kowalski CH, Kerkaert JD, Liu KW, Bond MC, Hartmann R, Nadell CD, Stajich JE, Cramer RA. 2019. Fungal biofilm morphology impacts hypoxia fitness and disease progression. Nat Microbiol 4:2430–2441.

9. Kowalski CH, Morelli KA, Schultz D, Nadell CD, Cramer RA. 2020. Fungal biofilm architecture produces hypoxic microenvironments that drive antifungal resistance. Proc Natl Acad Sci U S A 117:22473–22483.

10. Mowat E, Butcher J, Lang S, Williams C, Ramage G. 2007. Development of a simple model for studying the effects of antifungal agents on multicellular communities of Aspergillus fumigatus. J Med Microbiol 56:1205–1212.

11. Rajendran R, Williams C, Lappin DF, Millington O, Martins M, Ramage G. 2013. Extracellular DNA release acts as an antifungal resistance mechanism in mature Aspergillus fumigatus biofilms. Eukaryot Cell 12:420–429.

12. Rajendran R, Mowat E, McCulloch E, Lappin DF, Jones B, Lang S, Majithiya JB, Warn P, Williams C, Ramage G. 2011. Azole resistance of Aspergillus fumigatus biofilms is partly associated with efflux pump activity. Antimicrob Agents Chemother 55:2092–2097.

13. Wall G, Montelongo-Jauregui D, Vidal Bonifacio B, Lopez-Ribot JL, Uppuluri P. 2019. Candida albicans biofilm growth and dispersal: contributions to pathogenesis. Curr Opin Microbiol 52:1–6.

14. Nobile CJ, Fox EP, Nett JE, Sorrells TR, Mitrovich QM, Hernday AD, Tuch BB, Andes DR, Johnson AD. 2012. A recently evolved transcriptional network controls biofilm development in Candida albicans. Cell 148:126–138.

15. Kean R, Delaney C, Rajendran R, Sherry L, Metcalfe R, Thomas R, McLean W, Williams C, Ramage G. 2018. Gaining insights from Candida biofilm heterogeneity: One size does not fit all. Journal of Fungi 4.

16. Bonhomme J, Chauvel M, Goyard S, Roux P, Rossignol T, D’Enfert C. 2011. Contribution of the glycolytic flux and hypoxia adaptation to efficient biofilm formation by Candida albicans. Mol Microbiol 80:995–1013.

17. Kordana N, Johnson A, Quinn K, Obar JJ, Cramer RA. 2025. Recent developments in Aspergillus fumigatus research: diversity, drugs, and disease. Microbiology and Molecular Biology Reviews 89.

18. Latgé JP, Chamilos G. 2019. Aspergillus fumigatus and Aspergillosis in 2019. Clin Microbiol Rev 33.

19. Loussert C, Schmitt C, Prevost MC, Balloy V, Fadel E, Philippe B, Kauffmann-Lacroix C, Latgé JP, Beauvais A. 2010. In vivo biofilm composition of Aspergillus fumigatus. Cell Microbiol 12:405–410.

20. Beauvais A, Latgé J-P. 2015. Aspergillus Biofilm In Vitro and In Vivo. Microbiol Spectr 3.

21. Zarnowski R, Sanchez H, Covelli AS, Dominguez E, Jaromin A, Bernhardt J, Mitchell KF, Heiss C, Azadi P, Mitchell A, Andes DR. 2018. Candida albicans biofilm–induced vesicles confer drug resistance through matrix biogenesis. PLoS Biol 16:e2006872.

22. Gibbons JG, Beauvais A, Beau R, McGary KL, Latgé JP, Rokas A. 2012. Global transcriptome changes underlying colony growth in the opportunistic human pathogen Aspergillus fumigatus. Eukaryot Cell 11:68–78.

23. Hillmann F, Shekhova E, Kniemeyer O. 2015. Insights into the cellular responses to hypoxia in filamentous fungi. Curr Genet 61:441–455.

24. Losada L, Barker BM, Pakala SS, Pakala SS, Joardar V, Zafar N, Mounaud S, Fedorova N, Nierman WC, Cramer RA. 2014. Large-Scale Transcriptional Response to Hypoxia in Aspergillus fumigatus Observed Using RNAseq Identifies a Novel Hypoxia Regulated ncRNA. Mycopathologia 178:331–339.

25. Barker BM, Kroll K, Vödisch M, Mazurie A, Kniemeyer O, Cramer RA. 2012. Transcriptomic and proteomic analyses of the Aspergillus fumigatus hypoxia response using an oxygen-controlled fermenter. BMC Genomics 13:62.

26. Kroll K, Pähtz V, Hillmann F, Vaknin Y, Schmidt-Heck W, Roth M, Jacobsen ID, Osherov N, Brakhage AA, Kniemeyer O. 2014. Identification of hypoxia-inducible target genes of Aspergillus fumigatus by transcriptome analysis reveals cellular respiration as an important contributor to hypoxic survival. Eukaryot Cell 13:1241–1253.

27. Langfelder P, Horvath S. 2008. WGCNA: An R package for weighted correlation network analysis. BMC Bioinformatics 9.

28. Langfelder P, Horvath S. 2012. Fast R Functions for Robust Correlations and Hierarchical Clustering. J Stat Softw 46:1–17.

29. Caffrey-carr AK, Kowalski CH, Beattie SR, Blaseg NA, Upshaw CR, Thammahong A, Lust HE, Tang Y, Hohl TM, Cramer RA, Obar JJ. 2017. Interleukin 1alpha Is Critical for Resistance against Highly Virulent Aspergillus fumigatus Isolates 85:1–19.

30. Kowalski CH, Beattie SR, Fuller KK, McGurk EA, Tang YW, Hohl TM, Obar JJ, Cramer RA. 2016. Heterogeneity among isolates reveals that fitness in low oxygen correlates with Aspergillus fumigatus virulence. mBio 7:1–13.

31. Rizzetto L, Giovannini G, Bromley M, Bowyer P, Romani L, Cavalieri D. 2013. Strain Dependent Variation of Immune Responses to A. fumigatus: Definition of Pathogenic Species. PLoS One 8:e56651.

32. Sheppard DC, Doedt T, Chiang LY, Kim HS, Chen D, Nierman WC, Filler SG. 2005. The Aspergillus fumigatus StuA Protein Governs the Up-Regulation of a Discrete Transcriptional Program during the Acquisition of Developmental Competence. Mol Biol Cell 16:5866.

33. Twumasi-Boateng K, Yu Y, Chen D, Gravelat FN, Nierman WC, Sheppard DC. 2009. Transcriptional profiling identifies a role for BrlA in the response to nitrogen depletion and for StuA in the regulation of secondary metabolite clusters in Aspergillus fumigatus. Eukaryot Cell 8:104–115.

34. Bultman KM, Kowalski CH, Cramer RA. 2016. Aspergillus fumigatus virulence through the lens of transcription factors. Med Mycol 55:myw120.

35. Rittenour WR, Chen M, Cahoon EB, Harris SD. 2011. Control of Glucosylceramide Production and Morphogenesis by the Bar1 Ceramide Synthase in Fusarium graminearum. PLoS One 6:e19385.

36. Li S, Du L, Yuen G, Harris SD. 2006. Distinct ceramide synthases regulate polarized growth in the filamentous fungus Aspergillus nidulans. Mol Biol Cell 17:1218–1227.

37. Abramson J, Adler J, Dunger J, Evans R, Green T, Pritzel A, Ronneberger O, Willmore L, Ballard AJ, Bambrick J, Bodenstein SW, Evans DA, Hung CC, O’Neill M, Reiman D, Tunyasuvunakool K, Wu Z, Žemgulytė A, Arvaniti E, Beattie C, Bertolli O, Bridgland A, Cherepanov A, Congreve M, Cowen-Rivers AI, Cowie A, Figurnov M, Fuchs FB, Gladman H, Jain R, Khan YA, Low CMR, Perlin K, Potapenko A, Savy P, Singh S, Stecula A, Thillaisundaram A, Tong C, Yakneen S, Zhong ED, Zielinski M, Žídek A, Bapst V, Kohli P, Jaderberg M, Hassabis D, Jumper JM. 2024. Accurate structure prediction of biomolecular interactions with AlphaFold 3. Nature 630:493–500.

38. Varadi M, Bertoni D, Magana P, Paramval U, Pidruchna I, Radhakrishnan M, Tsenkov M, Nair S, Mirdita M, Yeo J, Kovalevskiy O, Tunyasuvunakool K, Laydon A, Žídek A, Tomlinson H, Hariharan D, Abrahamson J, Green T, Jumper J, Birney E, Steinegger M, Hassabis D, Velankar S. 2024. AlphaFold Protein Structure Database in 2024: providing structure coverage for over 214 million protein sequences. Nucleic Acids Res 52:D368–D375.

39. Jumper J, Evans R, Pritzel A, Green T, Figurnov M, Ronneberger O, Tunyasuvunakool K, Bates R, Žídek A, Potapenko A, Bridgland A, Meyer C, Kohl SAA, Ballard AJ, Cowie A, Romera-Paredes B, Nikolov S, Jain R, Adler J, Back T, Petersen S, Reiman D, Clancy E, Zielinski M, Steinegger M, Pacholska M, Berghammer T, Bodenstein S, Silver D, Vinyals O, Senior AW, Kavukcuoglu K, Kohli P, Hassabis D. 2021. Highly accurate protein structure prediction with AlphaFold. Nature 2021 596:7873 596:583–589.

40. Meng EC, Goddard TD, Pettersen EF, Couch GS, Pearson ZJ, Morris JH, Ferrin TE. 2023. UCSF ChimeraX: Tools for structure building and analysis. Protein Science 32:e4792.

41. Furukawa T, van Rhijn N, Chown H, Rhodes J, Alfuraiji N, Fortune-Grant R, Bignell E, Fisher MC, Bromley M. 2022. Exploring a novel genomic safe-haven site in the human pathogenic mould Aspergillus fumigatus. Fungal Genetics and Biology 161:103702.

42. Perdoni F, Signorelli P, Cirasola D, Caretti A, Galimberti V, Biggiogera M, Gasco P, Musicanti C, Morace G, Borghi E. 2015. Antifungal activity of Myriocin on clinically relevant Aspergillus fumigatus strains producing biofilm. BMC Microbiol 15:1–8.

43. Del Poeta M, Nimrichter L, Rodrigues ML, Luberto C. 2014. Synthesis and Biological Properties of Fungal Glucosylceramide. PLoS Pathog 10:e1003832.

44. Bertuzzi M, van Rhijn N, Krappmann S, Bowyer P, Bromley MJ, Bignell EM. 2021. On the lineage of Aspergillus fumigatus isolates in common laboratory use. Med Mycol 59:7–13.

45. Lofgren LA, Ross BS, Cramer RA, Stajich JE. 2022. Combined Pan-, Population-, and Phylo-Genomic Analysis of Aspergillus fumigatus Reveals Population Structure and Lineage-Specific Diversity. bioRxiv 2021.12.12.472145.

46. Barber AE, Sae-Ong T, Kang K, Seelbinder B, Li J, Walther G, Panagiotou G, Kurzai O. 2021. Aspergillus fumigatus pan-genome analysis identifies genetic variants associated with human infection. Nat Microbiol 6:1526–1536.

47. Horta MAC, Steenwyk JL, Mead ME, Braz dos Santos LH, Zhao S, Gibbons JG, Marcet-Houben M, Gabaldón T, Rokas A, Goldman GH. 2022. Examination of Genome-Wide Ortholog Variation in Clinical and Environmental Isolates of the Fungal Pathogen Aspergillus fumigatus. mBio 13.

48. Singh A, Wang H, Silva LC, Na C, Prieto M, Futerman AH, Luberto C, Del Poeta M. 2012. Methylation of glycosylated sphingolipid modulates membrane lipid topography and pathogenicity of Cryptococcus neoformans. Cell Microbiol 14:500–516.

49. Sentelle RD, Senkal CE, Jiang W, Ponnusamy S, Gencer S, Panneer Selvam S, Ramshesh VK, Peterson YK, Lemasters JJ, Szulc ZM, Bielawski J, Ogretmen B. 2012. Ceramide targets autophagosomes to mitochondria and induces lethal mitophagy. Nature Chemical Biology 2012 8:10 8:831–838.

50. Lingo DE, Shukla N, Osmani AH, Osmani SA. 2021. Aspergillus nidulans biofilm formation modifies cellular architecture and enables light-activated autophagy. Mol Biol Cell 32:1181–1192.

51. Rajasenan S, Osmani AH, Osmani SA. 2022. Modulation of sensitivity to gaseous signaling by sterol-regulatory hypoxic transcription factors in Aspergillus nidulans biofilm cells. Fungal Genetics and Biology 163:103739.

52. Kerkaert JD, Le Mauff F, Wucher BR, Beattie SR, Vesely EM, Sheppard DC, Nadell CD, Cramer RA. 2022. An Alanine Aminotransferase Is Required for Biofilm-Specific Resistance of Aspergillus fumigatus to Echinocandin Treatment. mBio 13.

53. Ewels P, Magnusson M, Lundin S, Käller M. 2016. MultiQC: summarize analysis results for multiple tools and samples in a single report. Bioinformatics 32:3047–3048.

54. Martin M. 2011. Cutadapt removes adapter sequences from high-throughput sequencing reads. EMBnet J 17:10–12.

55. Dobin A, Davis CA, Schlesinger F, Drenkow J, Zaleski C, Jha S, Batut P, Chaisson M, Gingeras TR. 2013. STAR: ultrafast universal RNA-seq aligner. Bioinformatics 29:15–21.

56. Anders S, Pyl PT, Huber W. 2015. HTSeq-A Python framework to work with high-throughput sequencing data. Bioinformatics 31:166–169.

57. Love MI, Huber W, Anders S. 2014. Moderated estimation of fold change and dispersion for RNA-seq data with DESeq2. Genome Biol 15:1–21.

58. Robinson MD, McCarthy DJ, Smyth GK. 2010. edgeR: a Bioconductor package for differential expression analysis of digital gene expression data. Bioinformatics 26:139–140.

59. Ritchie ME, Phipson B, Wu D, Hu Y, Law CW, Shi W, Smyth GK. 2015. limma powers differential expression analyses for RNA-sequencing and microarray studies. Nucleic Acids Res 43:e47–e47.

60. Gu Z. 2022. Complex heatmap visualization. iMeta 1:e43.

61. Horvath S, Langfelder P. 2011. Tutorials for the WGCNA package for R : WGCNA Background and glossary. Gene 7–9.

62. Garcia Lopez A, Albrecht-Eckardt D, Panagiotou G, Schäuble S. 2024. FungiFun3: systemic gene set enrichment analysis for fungal species. Bioinformatics 40.

63. Al Abdallah Q, Ge W, Fortwendel JR. 2017. A Simple and Universal System for Gene Manipulation in Aspergillus fumigatus: In Vitro -Assembled Cas9-Guide RNA Ribonucleoproteins Coupled with Microhomology Repair Templates. mSphere 2.

64. Schindelin J, Arganda-Carreras I, Frise E, Kaynig V, Longair M, Pietzsch T, Preibisch S, Rueden C, Saalfeld S, Schmid B, Tinevez J-Y, White DJ, Hartenstein V, Eliceiri K, Tomancak P, Cardona A. 2012. Fiji: an open-source platform for biological-image analysis. Nat Methods 9:676–82.

65. Hartmann R, Jeckel H, Jelli E, Singh PK, Vaidya S, Bayer M, Rode DKH, Vidakovic L, Díaz-Pascual F, Fong JCN, Dragoš A, Lamprecht O, Thöming JG, Netter N, Häussler S, Nadell CD, Sourjik V, Kovács ÁT, Yildiz FH, Drescher K. 2021. Quantitative image analysis of microbial communities with BiofilmQ. Nat Microbiol 6:151–156.

66. Beattie SR, Mark KMK, Thammahong A, Ries LNA, Dhingra S, Caffrey-Carr AK, Cheng C, Black CC, Bowyer P, Bromley MJ, Obar JJ, Goldman GH, Cramer RA. 2017. Filamentous fungal carbon catabolite repression supports metabolic plasticity and stress responses essential for disease progression. PLoS Pathog 13:1–29.

67. Edgar R, Domrachev M, Lash AE. 2002. Gene Expression Omnibus: NCBI gene expression and hybridization array data repository. Nucleic Acids Res 30:207–210.

